# SAFA facilitates chromatin opening of immune genes through interacting with nascent antiviral RNAs

**DOI:** 10.1101/2021.07.06.451336

**Authors:** Lili Cao, Yunfei Li, Yujie Luo, Xuefei Guo, Shengde Liu, Siji Li, Junhong Li, Zeming Zhang, Yingchi Zhao, Qiao Zhang, Feng Gao, Xiong Ji, Yiguang Wang, Xiang Gao, Fuping You

**Affiliations:** Institute of Systems Biomedicine, Department of Immunology, School of Basic Medical Sciences, Beijing Key Laboratory of Tumor Systems Biology, Peking University Health Science Center, Beijing, 100191, China; Department of Pharmaceutics, School of Pharmaceutical Sciences, Peking University, Beijing 100191, China; University of Chinese Academy of Sciences, CAS Key Laboratory of Infection and Immunity, National Laboratory of Macromolecules, Institute of Biophysics, Chinese Academy of Sciences, Beijing, 100101, China; School of Medicine, Jinan University, Guangzhou, Guangdong, 510632, China; Key Laboratory of Cell Proliferation and Differentiation of the Ministry of Education, School of Life Sciences, Peking-Tsinghua Center for Life Sciences, Peking University, Beijing, 100871, China; State Key Laboratory of Microbial Technology, Microbial Technology Institute, School of life science, Shandong University, Qingdao, 266000, China

**Author notes:** These authors contributed equally (Lili Cao, Yunfei Li, Yujie Luo). Corresponding author: Fuping You Ph.D, Institute of Systems Biomedicine, Department of Immunology, Beijing Key Laboratory of Tumor Systems Biology, Peking University Health Science Center, 100191, Beijing, China.

## Abstract

Regulation of chromatin accessibility determines the transcription activities of genes, which endow the host with function-specific gene expression patterns. It remains unclear how chromatin accessibility is specifically directed, particularly, during host defense against viral infection. We previously reported that the nuclear matrix protein SAFA surveils viral RNA and regulates antiviral immune genes expression. However, how SAFA regulates the expression and what determines the specificity of antiviral immune genes remains unknown. Here, we identified that the depletion of SAFA specifically decreased the chromatin accessibility, activation and expression of virus induced genes in a genome-wide scale after VSV infection. SAFA exclusively bound with antiviral related RNAs, which mediated the specific opening of the according chromatin and robust transcription of these genes. Knockdown of these associated RNAs dampened the accessibility of corresponding genes in an extranuclear signaling pathway dependent manner. Moreover, VSV infection cleaved SAFA protein at the C-terminus which deprived its RNA binding ability for immune evasion. Thus, our results demonstrated that SAFA and the interacting RNA products during viral infection collaborate and remodel chromatin accessibility to facilitate antiviral innate immune response.

## Introduction

In the eukaryotic cell nucleus, the chromatin structures are hierarchical ordered, ranging from kilobase to megabase scales (Belmont, 2014; Bonev and Cavalli, 2016). The multiple levels including nucleosome, loops, topologically associated domains (TADs), A/B compartments and territories (Cremer and Cremer, 2010; Lieberman-Aiden et al., 2009; Pope et al., 2014). Chromatin is the template of all DNA-related processes. The proper regulation of chromatin structure and the subsequent accessibility of DNA are essential for the performance of numerous cellular functions (Agarwal and Rao, 1998; Hao et al., 2019; Kim and Kaang, 2017; Masliah-Planchon et al., 2015; Nguyen et al., 2001). Upon viral infection, the innate immune response provides a first line of defense, allowing rapid production of variegated anti-viral cytokines (Akira and Takeda, 2004; Beutler, 2004; Takeuchi and Akira, 2010). This process is primarily controlled by dynamic organization of the genome, which reprogrammed the specific genomic regions from a condensed state to a transcriptionally accessible state (Klemm et al., 2019; Lanctot et al., 2007). Hence, there should be a precise molecular mechanism underpinning the reprogramming of defensive responses.

Processes involved in the alteration of chromatin accessibility are diverse, including post-translational modifications of histones, incorporation of histone variants, DNA methylation and ATP-dependent chromatin remodeling (Bao et al., 2019; Deuring et al., 2000; Govin et al., 2004; Venkatesh and Workman, 2015). There is accumulating evidence indicating that RNAs also play an important role (Caudron-Herger and Rippe, 2012; Dong et al., 2020; Gupta et al., 2010; Han and Chang, 2015; Mousavi et al., 2013). The RNA encoded by HOXC locus represses transcription of the HOXD locus through interacting with the polycomb repressive complex 2 (PRC2) (Rinn et al., 2007). During the process of mammal X-chromosome inactivation, the stable repression of all X-linked genes is mediated by the long noncoding RNA, Xist, which is transcribed from specific X-linked sequences (Gendrel and Heard, 2014). Xist induces a cascade of chromatin changes, including post-translational histone modifications and DNA methylation, by interacting with multiple proteins. These findings implicate the regulation roles of RNAs in gene expression are not only broad spectrum but also related to corresponding locus. 62%–75% of the human genome is capable of producing various RNA species, but less than 2% encodes proteins (Djebali et al., 2012). RNAs reflect the direct production of the genetic information encoded by genomes. In addition, RNAs production is highly dynamic that different species and amounts of RNAs are produced at different stages of transcription (Morris and Mattick, 2014; Roden and Gladfelter, 2021). These led us to wonder whether determination and characterization of the regulatory regions of chromatin are regulated by the RNA product during viral infection.

Scaffold attachment factor A (SAF-A), also known as heterogeneous ribonucleoprotein U (HNRNP-U), is an abundant nuclear matrix associated protein (Fackelmayer et al., 1994). Traditionally, SAFA is an RNA-binding protein mainly involved in regulating gene transcription and RNA splicing (Geuens et al., 2016). Several reports suggest that SAFA plays a critical role in the recruitment of Xist RNA in inactive X chromosome (Kolpa et al., 2016). Recently, SAFA was demonstrated to play a central role in regulating chromatin architecture. The *in situ* Hi-C assay showed that SAFA mainly binds to active chromatin (Fan et al., 2018). Disruption of SAFA leads to compartment switching from B to A and reduces the TAD boundary strengths at borders between two types of compartments (Fan et al., 2018). Nozawa et al. reported that oligomerized SAFA remodels interphase chromatin structures through interaction with nascent RNAs (Nozawa et al., 2017). SAFA oligomerization decompacts large-scale chromatin structure while SAFA deficiency or monomerization promotes aberrant chromosome folding (Nozawa et al., 2017). Our previous study suggests that SAFA surveils viral RNA in the nucleus and facilitates innate immune response by activating antiviral enhancers and super-enhancers (Cao et al., 2019a). Interestingly, this process was also dependent on SAFA oligomerization. Viral infection induces SAFA oligomerization, which is essential for the activation of antiviral immune responses (Cao et al., 2019a). However, it is unknown if or how SAFA regulates the accessibility of the specific chromatin locus coding antiviral genes during virus infection.

In the present study, by combining Assay for Transposase-Accessible Chromatin with high throughput sequencing (ATAC-seq), Chromatin immunoprecipitation followed by sequencing (ChIP-seq) and bulk RNA-sequencing (RNA-seq), we assessed the genome-wide chromatin accessibility and gene expression in wild type and SAFA deficient cells after viral infection, and found that SAFA was essential for the chromatin accessibility and activation of antiviral immune genes. In addition, this process is dependent on the association of SAFA with nascent transcripts. Mechanistically, RNAs produced during viral infection interacted with SAFA and mediated the accessibility of related chromatin regions in an extranuclear antiviral signaling pathways dependent manner, which mediated the expression of antiviral genes. Intriguingly, on the other hand, viral infection induced cleavage of SAFA that separated its RNA binding domain for immune evasion. Hence, the canonical antiviral pathways directed production of nascent antiviral transcripts, which bound to and activated SAFA, and in turn SAFA further facilitated transcription of these antiviral genes by increasing the openness of chromatin.

## Results

### SAFA deficiency decreased the chromatin accessibility of antiviral immune genes

To explore the potential role of SAFA in regulating chromatin accessibility during viral infection, we performed ATAC-seq and RNA-seq analysis in wild type and SAFA deficient (*SAFA*^-/-^) THP-1 cells (Figure S1A). SAFA deficiency leaded to an extensive decrease in chromatin accessibility at both the promoter and the UTR regions during VSV infection (Figure 1A and S1B). Intriguingly, the decrease of chromatin accessibility by SAFA disruption exclusively took place at the locus governing the expression of viral induced genes, but not housekeeping locus (Figure 1B and S1C). The openness of the locus induced over 1000 fold in THP-1 cells after VSV infection was apparently impaired due to SAFA depletion, while the locus where the accessibility had no obvious impact during viral infection also showed no significant differences in infected *SAFA*^-/-^ cells (Figure 1B and S1C). The Gene Ontology (GO) term enrichment analysis showed that these genes significantly affected by SAFA depletion were involved in type I interferon signaling pathway and host defense response to virus (Figure 1C). Type I IFNs and ISGs are potent innate antiviral immune response effectors. Consistently, the chromatin accessibility of related ISGs were greatly decreased in SAFA deficient cells (Figure 1D, 1E and S1D). *CXCL10, CCL5, DDX58, ISG15, MX1* and *OASL* are known to code important antiviral effectors. *CXCL10, ISG15, MX1* and *OASL* are important interferon-stimulated genes (ISGs) (Schneider et al., 2014; Schoggins and Rice, 2011). CCL5 is a T cell chemoattractant that is critical for immune control of viral infections (Crawford et al., 2011). Retinoic acid-inducible gene 1 (RIG-I), which is encoded by *DDX58* gene, is critical for sensing of cytoplasmic viral RNA to initiate and modulate antiviral innate immunity (Loo and Gale, 2011). The virus induced chromatin accessibility of these genes was robustly decreased in *SAFA*^-/-^ cells (Figure 1E). The transcription factor enrichment analysis revealed a loss of accessibility for genes with IRF3, IRF1, IRF8 and IRF2 motifs (Figure 1F). Notably, interferon regulatory factors (IRF) target genes have a critical role in the regulation of host defense (Taniguchi et al., 2001).

**Figure 1with figure supplement 1- source data 1.**
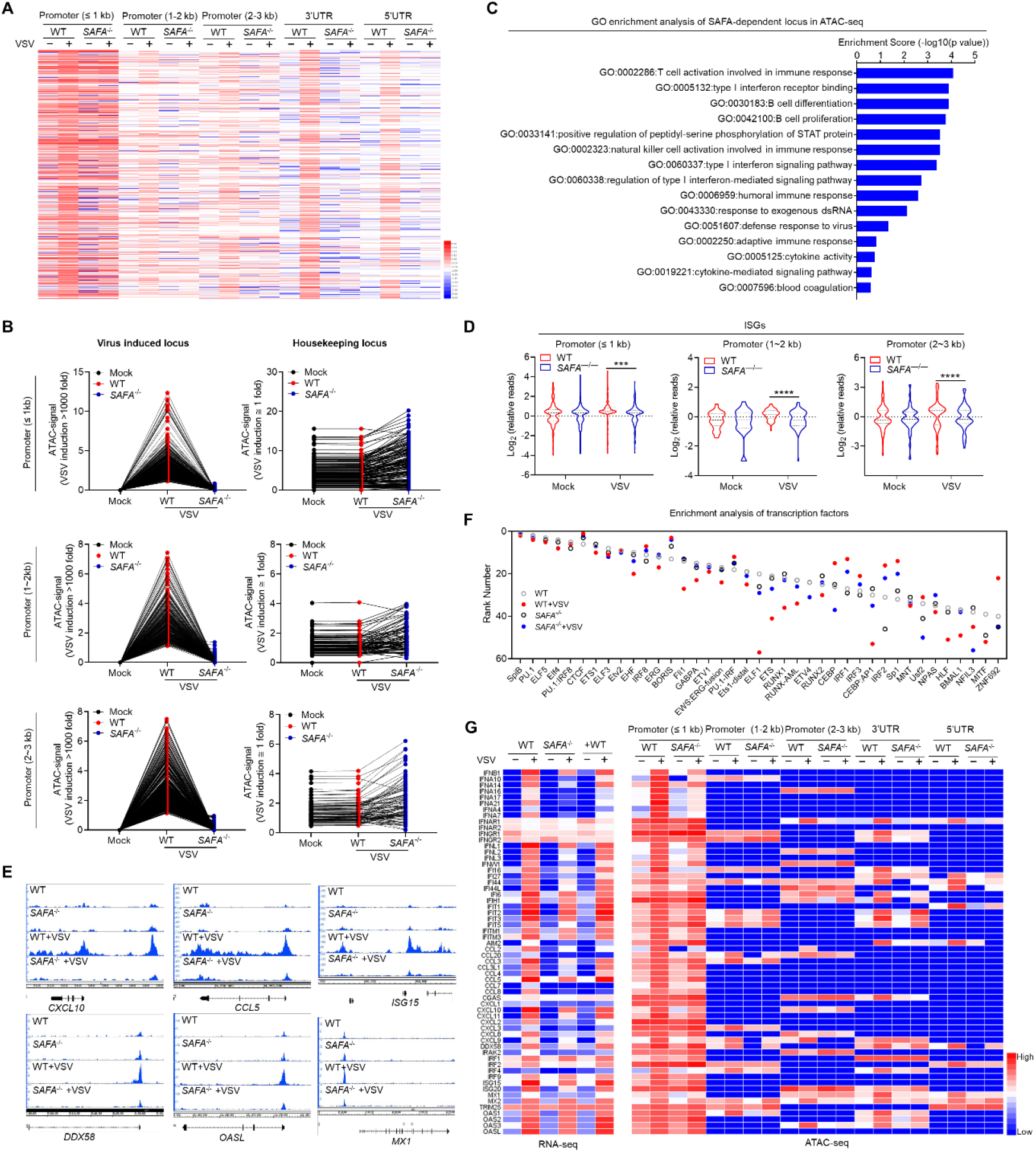
SAFA deficiency decreased the chromatin accessibility of antiviral immune genes. (A) Heatmap showing the ATAC-seq signal in Wild-type (WT) and *SAFA*^-/-^ THP-1 cells with VSV infection for 6 hours. (B) Line graph showing SAFA in regulation of VSV-induced accessible locus and housekeeping locus in ATAC-seq. (C) GO term enrichment analysis of genes significantly affected by SAFA depletion in ATAC-seq. (D) Violin graph showing ISGs affected by SAFA depletion in ATAC-seq. (E) Genome browser views of ATAC-seq signal for the indicated genes. (F) Transcription factor enrichment analysis of ATAC-seq. (G) Heatmap comparing ATAC-seq signal and RNA-seq signal of indicated genes. \*\*\**p* < 0.001, \*\*\*\**p* < 0.0001 (Student’s t test; D). Data were pooled from two independent experiments (A-D, F and G). Data were representative of two independent experiments (E).

Integrated analysis of RNA-seq and ATAC-seq revealed that genes with reduced chromatin accessibility also showed significant lower expression levels (Figure 1G). Correspondingly, the downregulated genes in SAFA deficient cells after VSV infection were mainly enriched in innate immune response to virus (Figure S1E). Together, these results suggest that SAFA mediates the chromatin accessibility of antiviral immune genes after viral infection.

### SAFA deficiency decreased the activation of antiviral immune genes

Enhancers and promoters are key regulatory DNA elements that control gene expression (Juven-Gershon and Kadonaga, 2010; Ong and Corces, 2011). The accessible chromatin reflects a permission for physical interactions of transcription machineries with enhancers and promoters, which regulate transcriptional activation (Klemm et al., 2019). To confirm the role of SAFA in enhancing the chromatin accessibility of antiviral genes, we performed ChIP-seq analysis with H3 lysine 27 acetylation (H3K27ac) antibody. Enhancer activation was marked by high level of H3K27ac (Creyghton et al., 2010). Our previous results showed that SAFA facilitates distal enhancer activation of type I IFN (Cao et al., 2019a). Here we assessed the impact of SAFA on activation of virus-induced enhancers in a genome-wide scale (Figure 2A and S2A). SAFA deficiency downregulated the enhancer activation globally after VSV infection (Figure 2A). There were 24799 enhancers in resting wild type THP-1 cells, 27828 after VSV infection for 8 hours, and 28964 after VSV infection for 24 hours. As for *SAFA*^-/-^ cells, there were 24499 enhancers in resting cells, 26225 after VSV infection for 8 hours, and 28150 after VSV infection for 24 hours (Figure S2B). The consequences can be even greater at the early stage of infection (Figure S2C). Moreover, these enhancers inactivated by SAFA disruption were mainly involved in response to virus infection (Figure 2B).

**Figure 2 with figure supplement 2.**
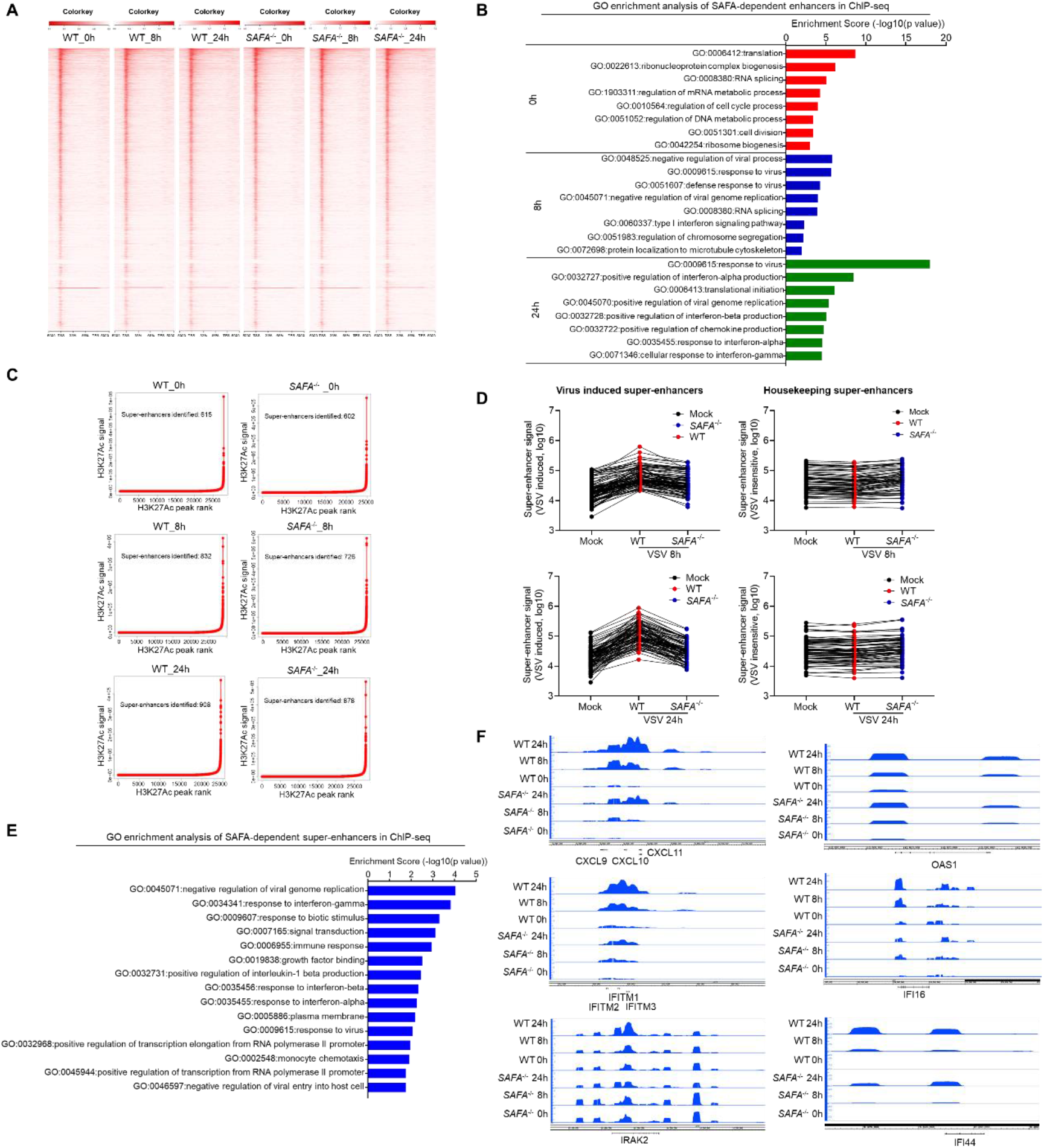
SAFA deficiency decreased the activation of antiviral immune genes. (A) Heatmap showing the ChIP-seq signal enrichment around the TSSs of H3K27ac in WT and *SAFA*^-/-^ THP-1 cells with VSV infection for 8 or 24 hours. (B) GO term enrichment analysis of enhancers-related genes affected by SAFA depletion in ChIP -seq. (C) Delineation of super-enhancers based on H3K27Ac occupancy in WT and *SAFA*^-/-^ THP-1 cells with VSV infection using the ROSE algorithm. (D) Line graph showing SAFA in regulation of VSV-induced and housekeeping supper-enhancer formation. (E) GO term enrichment analysis of super-enhancers related genes affected by SAFA depletion in ChIP -seq. (F) Genome browser views of ChIP -seq signal for the indicated genes. Data were pooled from two independent experiments (A-E). Data were representative of two independent experiments (F).

Super-enhancers are clusters of enhancers across a long range of genomic DNA, which drive expression of genes that define cell state (Hnisz et al., 2013; Pott and Lieb, 2015). It was also marked by H3K27ac. Further analysis showed that SAFA is required for the activation of super-enhancers induced by viral infection. There were 615 super-enhancers in resting THP-1 cells, 832 after VSV infection for 8 hours, and 908 after VSV infection for 24 hours. In *SAFA*^-/-^ cells, there were 602 super-enhancers in untreated cells, 726 after VSV infection for 8 hours, and 878 after VSV infection for 24 hours (Figure 2C). SAFA deficiency decreased the formation of super-enhancers after VSV infection, especially at the early stage of infection (Figure 2D and S2D). Meanwhile, SAFA depletion showed no obvious impact on the formation of super-enhancers that insensitive to VSV infection (Figure 2D), suggesting that SAFA mainly affected the induction of super-enhancers related to viral infection. The enrichment analysis suggested that these super-enhancers associated genes significantly downregulated by SAFA depletion were involved in immune responses and host defense to virus (Figure 2E). Consistently, the super-enhancer formation of *CXCL9/10/11, OAS1, IFITM1/2/3, IRAK2, IFI16* and *IFI44* genes was robustly decreased in SAFA mutant cells after virus infection (Figure 2F). *CXCL9/10/11*, *OAS1, IFITM1/2/3, IFI16* and *IFI44* are important ISGs (Schneider et al., 2014; Schoggins and Rice, 2011). *IRAK2* is an essential adaptor of the Toll-like receptors (TLR) signaling pathway, which is important for downstream defense molecules production (Meylan and Tschopp, 2008). Moreover, the majority of these impaired super-enhancer-driven genes were protein-coding genes, but there was also a considerable part of non-coding genes (Figure S2E). Therefore, SAFA is required for virus induced enhancers/super-enhancers activation.

### RNA binding activity of SAFA is critical for increasing the accessibility of anti-viral chromatin

We then investigated the mechanism by which SAFA specifically increased chromatin accessibility of antiviral genes after infection. In the interphase, SAFA remodels the chromatin structures through the interaction with nascent RNAs (Nozawa et al., 2017). There is an increasing body of evidence suggesting that RNAs are involved in regulation of chromatin accessibility. By using genome-wide binding profiling, Kambiz et al. showed that eRNAs regulate genomic accessibility of the transcriptional complex to defined regulatory regions (Mousavi et al., 2013). Dong et al. reported that the lncRNA, *LncMyoD*, regulates lineage determination and progression through modulating chromatin accessibility (Dong et al., 2020). Our previous results suggest that SAFA facilitates anti-viral innate immune responses, which is also dependent on the RNA-binding ability (Cao et al., 2019a). These prompted us to investigate whether the regulation of chromatin accessibility by SAFA during virus infection is also dependent on the interaction with RNAs.

Structurally, SAFA contains an N-terminal DNA-binding domain, an ATP-binding AAA+ domain, a SPRY domain and an RNA-binding RGG repeat at the C-terminal (Erzberger and Berger, 2006; Romig et al., 1992). We rescued the full-length (Flag-SAFA) and RGG domain depleted (Flag-Del-RGG) Flag tagged SAFA plasmids into *SAFA*^-/-^ THP-1 cells, which enabled their stable expression (Figure S3A). Further, we did ATAC-seq and RNA-seq in both cell lines (Figure 3A). The results showed that after VSV infection, the genome-wide chromatin accessibility was downregulated in RGG domain mutated cells compared with that in Flag-SAFA cells (Figure 3B and S3B). The GO enrichment analysis showed that these downregulated genes were mainly involved in immune response and response to interferon (Figure 3C). Consistently, the chromatin accessibility of ISGs were significantly decreased in RGG domain depleted cells (Figure 3D, 3E and S3C), and the genome-wide transcription factor enrichment analysis inferred an impaired association of genes with IRF motifs in them (Figure 3F).

**Figure 3 with figure supplement 3 - source data 3.**
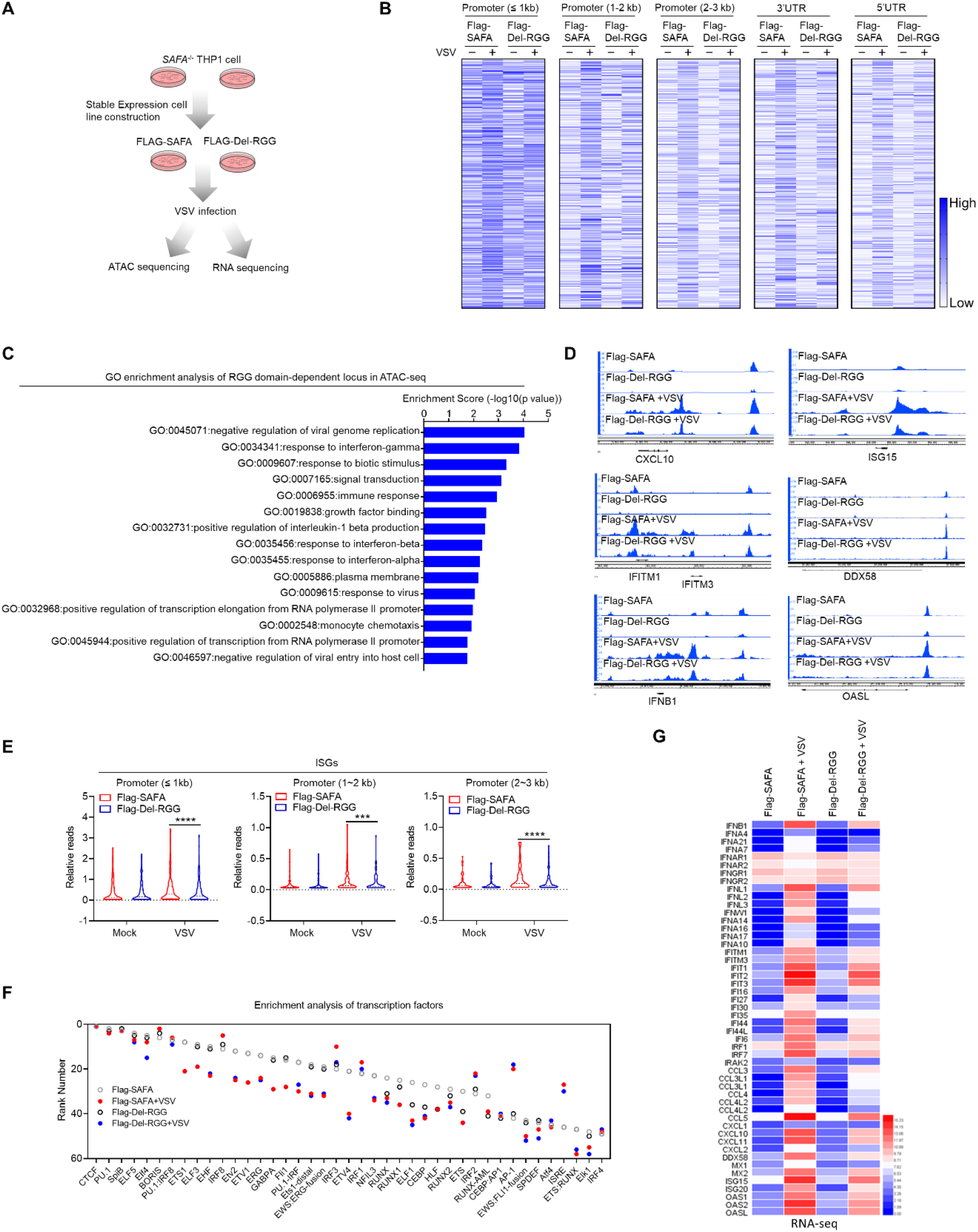
RNA binding activity of SAFA is critical for increasing the accessibility of anti-viral chromatin. (A) Models depicting the ATAC-seq and RNA-seq assay in Flag-SAFA and Flag-Del-RGG stably expressed *SAFA*^-/-^ THP-1 cells with VSV infection for 6 hours. (B) Heatmap showing the ATAC-seq signal. (C) GO term enrichment analysis of genes significantly affected by RGG domain depletion in ATAC-seq. (D) Genome browser views of ATAC-seq signal for the indicated genes. (E) Violin graph showing ISGs affected by RGG domain depletion in ATAC-seq. (F) Transcription factor enrichment analysis of ATAC-seq. (G) Heatmap showing RNA-seq signal for the indicated genes. \*\*\**p* < 0.001, \*\*\*\**p* < 0.0001 (Student’s t test; E). Data were pooled from two independent experiments (B, C and E-G). Data were representative of two independent experiments (D).

Moreover, the expression of anti-viral genes was obviously downregulated in Flag-Del-RGG cells (Figure 3G). RGG domain depletion mainly affected the regulation of type I interferon-mediated signaling pathway following viral infection (Figure S3D). These results suggest that the RNA-binding ability is essential for SAFA in maintaining chromatin accessibility of antiviral immune genes during viral infection.

### SAFA interacted with antiviral related RNAs in a time-dependent manner during viral infection

To gain further insight into the role of RNA-binding ability of SAFA in chromatin structure regulation after viral infection, we performed RNA immunoprecipitation sequencing (RIP-seq) of THP-1 cells following VSV infection for 6 hours and 24 hours (Figure S4) (Zhao et al., 2010). SAFA showed differential binding profiles at different stages of viral infection (Figure 4A). The RNAs interacting with SAFA were increased by 42.54% and 51.01% after VSV infection for 6 hours and 24 hours respectively (Figure 4B). More than 50% of the total increased SAFA binding RNAs were protein coding mRNAs after VSV infection (Figure 4B). There is also a considerable part of noncoding RNAs (ncRNAs), especially lncRNA (Figure 4B).

**Figure 4 with figure supplement 4.**
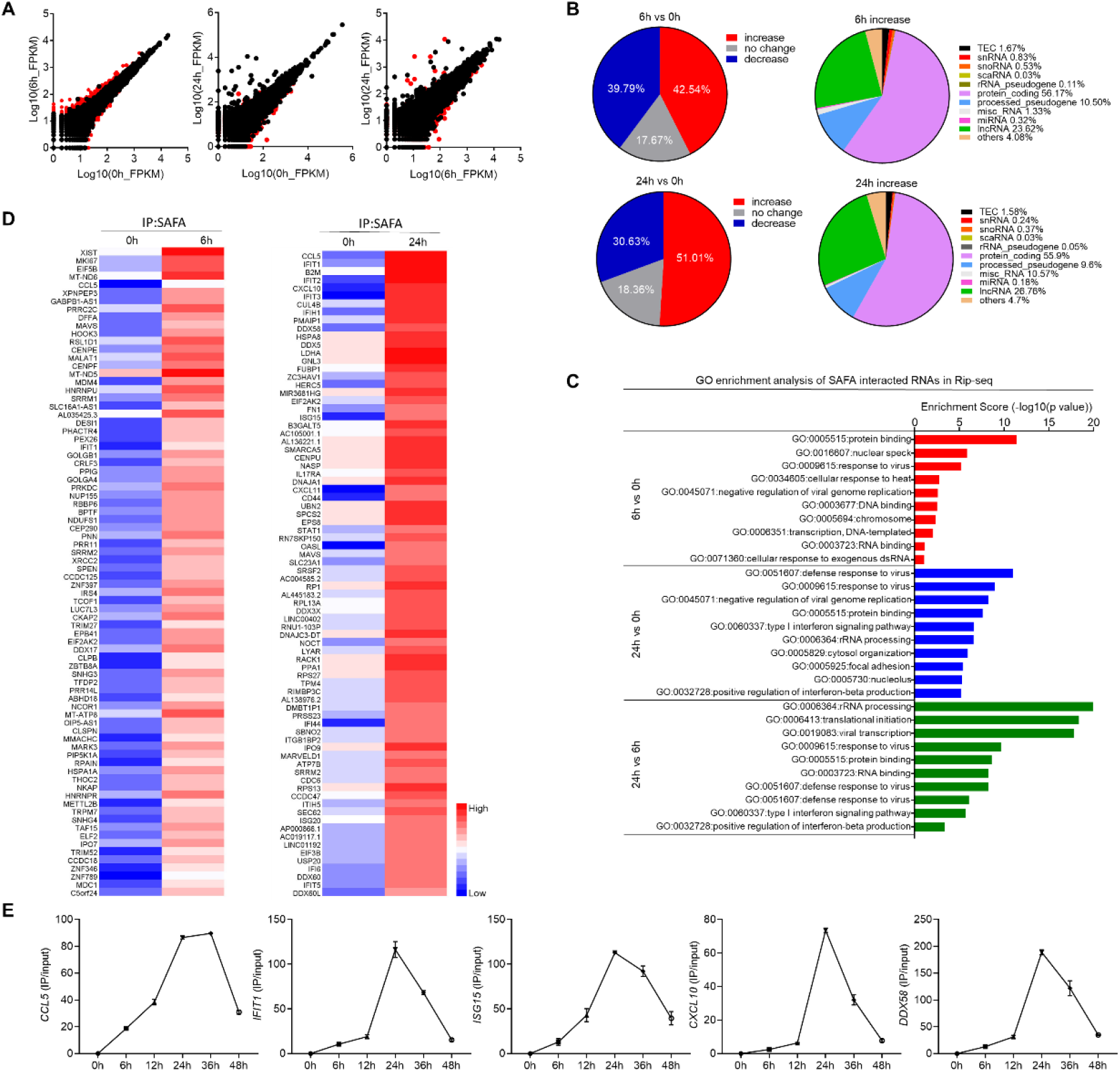
SAFA interacted with antiviral related RNAs in a time-dependent manner during viral infection. (A) Scatter diagram showing differential RNA binding profiles of SAFA in THP-1 cells with VSV infection for 6 hours or 24 hours. (B) Pie chart showing the changes of RNAs interacted with SAFA in RIP-seq upon VSV infection (left); pie chart showing the distribution profile of RNAs with increased interaction with SAFA after VSV infection (right). (C) GO term enrichment analysis of RNAs interacted with SAFA in RIP-seq. (D) Heatmap showing RIP-seq signal for the indicated RNAs. (E) Line graph showing time-dependent RNA binding manner of indicated genes with VSV infection for indicated times. Data were pooled from two independent experiments (A-D). Data were pooled from three experiments (E).

GO enrichment analysis revealed that these RNAs differentially interacting with SAFA were mainly involved in response to virus and protein binding after viral infection for 6 hours (Figure 4C). At the later stage of infection, these associated RNAs were almost entirely related to defense response to virus (Figure 4C). Among those RNAs, *CCL5*, *IFIT1/2/3, CXCL10, MAVS, DDX58, ISG15, OASL* and *IFI44* are known to encode important innate antiviral effectors (Figure 4D) (Crawford et al., 2011; Loo and Gale, 2011; Schneider et al., 2014; Schoggins and Rice, 2011). Furthermore, the results of more detailed time points suggested that the interaction of SAFA with these RNAs showed a time-dependent manner, in which the binding first increased with the time after VSV infection, peaked at about 24 hours and then dropped around 48 hours (Figure 4E). Thus, these results showed that SAFA interacted with antiviral related RNAs in a time dependent manner after viral infection.

### SAFA-interacting RNA mediated specific chromatin remodeling in an extranuclear pathway dependent manner

There is accumulating evidence suggests that RNA molecules are components of and play regulatory roles at different stages of transcription. Recent studies have shown that RNAs produced during early steps in transcription initiated the transcriptional condensate formation (Henninger et al., 2021). Moreover, the regulation roles of RNAs in gene expression showed locus-specific characteristics, which tends to regulate the expression of adjacent or related genes (Dong et al., 2020; Gendrel and Heard, 2014; Mousavi et al., 2013; Rinn et al., 2007). The RNA binding-dependent regulatory activity of SAFA, coupled with evidence that the associated RNAs are mainly antiviral innate immunity related, led us to wonder whether the SAFA-interacting RNA mapped or characterized the regulatory regions of accessible chromatin during viral infection.

To explore the potential role of SAFA-interacting RNA in regulating chromatin accessibility, we sought to knockdown the specific RNA product by CRISPR-Cas13d system and further detect the chromatin accessibility with ATAC-qPCR after viral infection (Figure 5A) (Kushawah et al., 2020). Results showed that this system could induce efficient RNA knockdown after VSV infection, and we selected CRISPR RNA (crRNA) 3# for *IFIT1*, crRNA 1# for *ISG15*, crRNA 2# for *CXCL10*, crRNA 3# for *CCL5*, crRNA 1# for *IFNB1* and crRNA 2# for *DDX58* for further experiments (Figure 5B). Interestingly, cells expressing specific crRNA were not able to sustain corresponding chromatin accessible during viral infection (Figure 5C). Consistently, the corresponding mRNA expression were apparently knockdown (Figure S5A). These results suggest that RNA product interacting with SAFA after viral infection mediated the accessibility of corresponding chromatin regions.

**Figure 5 with figure supplement 5.**
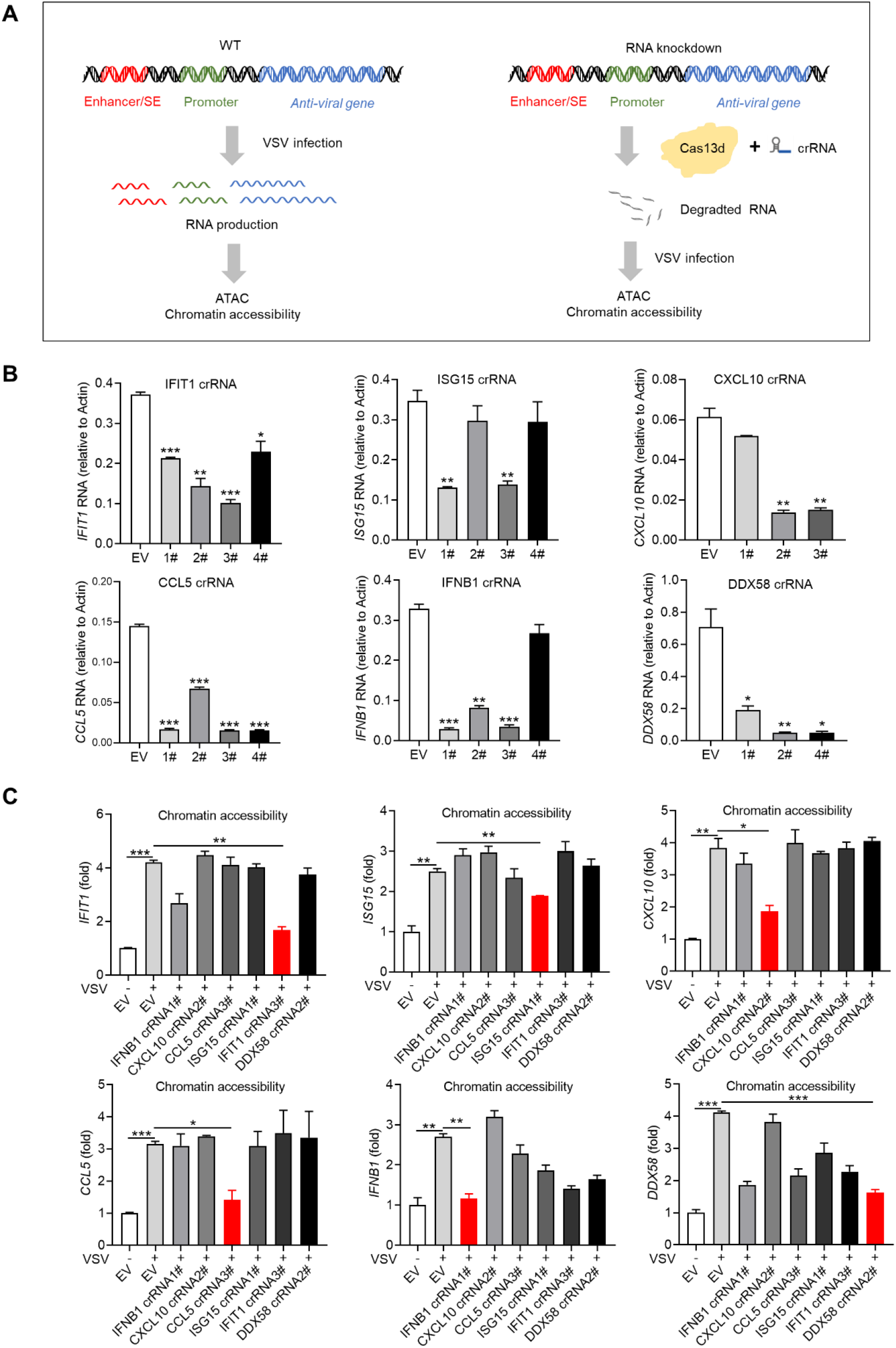
RNA product interacted with SAFA mediated specific chromatin remodeling during viral infection. (A) Models depicting the experiment design of knocking down RNA by CRISPR-Cas13d system and further detecting the chromatin accessibility with ATAC-qPCR after VSV infection. (B) Histogram showing the knockdown efficiency of crRNA of indicated RNAs after VSV infection for 18 hours. (C) ATAC-qPCR results showing the chromatin accessibility of indicated genes after the related RNA knockdown with or without VSV infection for 18 hours. Empty vector (EV) was used as control. \**p* < 0.05, \*\**p* < 0.01, \*\*\**p* < 0.001 (Student’s t test, B and C). Data were pooled from three independent experiments (B and C). Error bars, SEM. n = 3 cultures.

Notably, almost all of these VSV induced RNAs are known to require the RLR pathways for induction, and MAVS, a key adaptor protein of RLR signlaing, mediates the recruitment of downstream transcription factors NFκB and IRFs and the transcriptional activation of interferons and proinflammatory cytokine genes. We thus infected wild-type and *MAVS*^-/-^, *IRF3*^-/-^ THP-1 cells with VSV and did ATAC-qPCR. Compared with those in wild-type cells, the inducible accessibilities of *IFIT1, CXCL10, CCL5, IFNB1*, *DDX58* and *ISG15* were largely decreased in *MAVS*^-/-^ and *IRF3*^-/-^ cells (Figure S5B). The strong dependence of chromatin accessibility on MAVS and IRF3 supports that extranuclear signaling pathways confer a requirement for remodeling of chromatin after viral infection.

### Virus-mediated cleavage separates the RNA-binding domain from SAFA

Interestingly, in VSV infected THP-1 cells, we repeatedly observed a protein band just under the SAFA protein, with a smaller molecular weight of 10–20 kDa, and this phenotype appeared as early as 1 hour after VSV treatment (Figure 6A). Similar results were observed in primary murine bone marrow derived macrophages (BMDM) and HEK293T cells (Figure 6A and S6A). Thus, we reasoned that VSV infection might promote cleavage of SAFA protein.

**Figure 6 with figure supplement 6 - source data 6.**
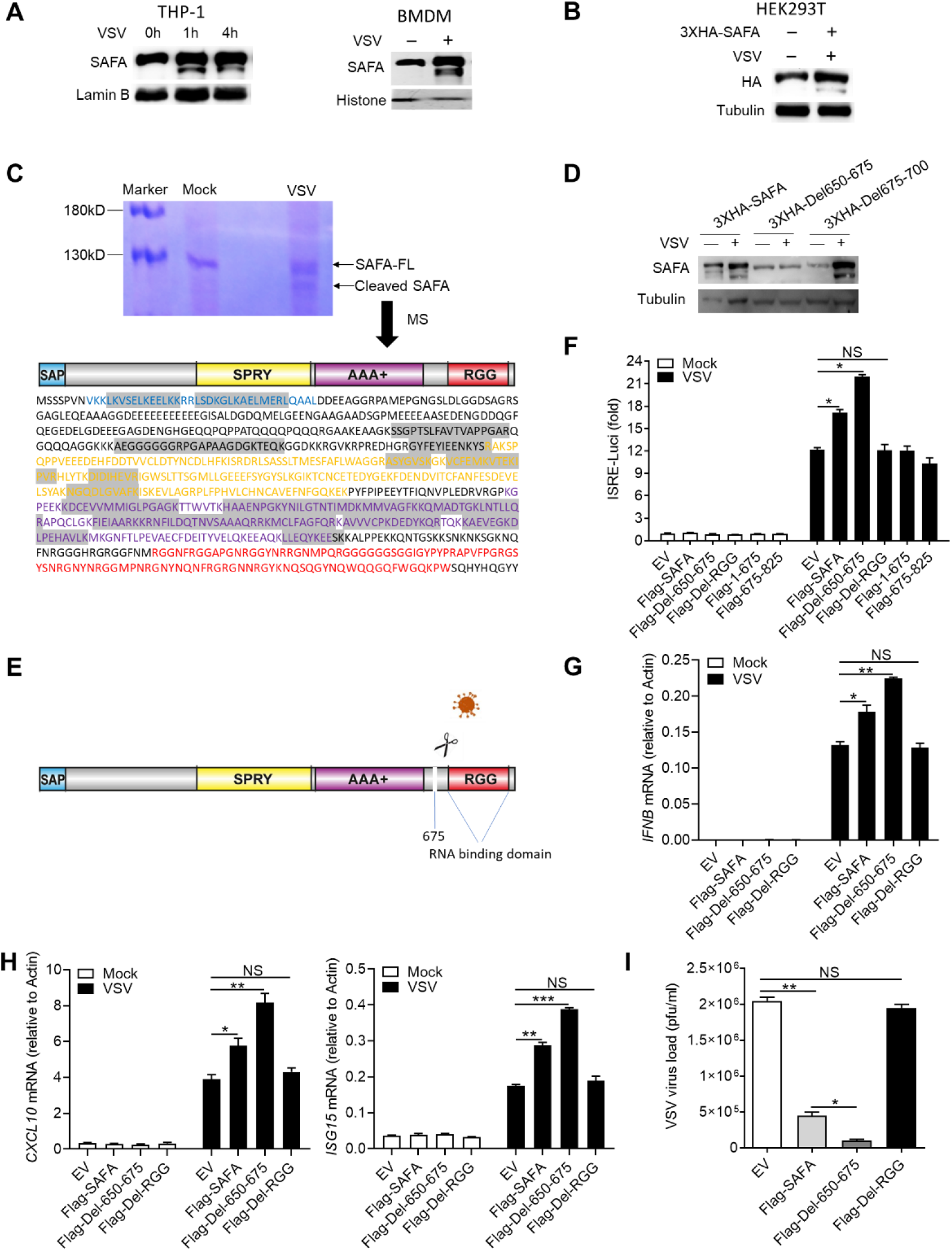
Virus-mediated cleavage of SAFA separates the RNA-binding domain. (A) Immunoblotting results showing the expression of indicated protein in THP-1 cells infected with VSV for indicated times or BMDM infected with VSV for 4 hours. (B) HEK293T cells were transfected with 3XHA-SAFA plasmids, and then infected with VSV for 4 hours followed by immunoblotting. (C) THP-1 cells were infected with VSV for 4 hours followed by immunoprecipitation and coomassie brilliant blue staining. The cleaved band was cut out for mass spectrum assay. The detected amino acid sequences were marked by grey background. (D) HEK293T cells were transfected with indicated plasmids, and then infected with VSV for 4 hours followed by immunoblotting. (E) Models depicting VSV infection induced cleavage of SAFA. (F) HEK293T cells were transfected with indicated plasmids before infection with VSV for 24 hours and then type I interferons in the supernatants were detected by bioassay. (G-I) THP-1 mutants generated by overexpressing indicated lentivirus plasmids were infected with VSV for 24 hours and the expression levels of *IFNB* (G) and *CXCL10, ISG15* (H) were detected by qPCR. The viral load was detected by plaque assay (I). \**P* < 0.05, \*\**P* < 0.01 and \*\*\**P* < 0.001 (Student’s t-test). Data were representative of three independent experiments (A-D). Data were pooled from 3 independent experiments (F-I). Error bars, SEM. n = 3 cultures.

To prove this hypothesis, we infected HEK293T cells that overexpressed N-terminal 3XHA-tagged SAFA with VSV, and a clear band similar to endogenous result was observed (Figure 6B). This result indicated that VSV infection led to a cut of SAFA at the C terminus, thus releasing the big N-terminal fragment. Further, we immunoprecipitated the cleaved band with SAFA antibody and visualized it with coomassie brilliant blue R250 staining. Then this band was cut out and sent for mass spectrometry analysis. The detected amino acid sequences located in the N-terminal SAP domain and the middle SPRY and AAA+ domain, but not the C-terminal RGG domain (Figure 6C), and the last amino acid detected was Lys^675^. However, when we infected cells that overexpressed Lys^675^ mutant with VSV, the cleaved bands still appeared (Data not shown). Further, we constructed two 3XHA-tagged deletion mutants that deleted the 650 to 675 amino acids (3XHA-Del650-675) or the 675-700 amino acids (3XHA-Del675-700). It was found that 3XHA-Del650-675 mutant showed resistance to VSV-infection induced cleavage of SAFA, indicating that amino acids 650 to 675 were VSV targeted sequences (Figure 6D). These results suggest that VSV infection mediated cleavage of SAFA which separates the RNA-binding domain (Figure 6E).

In agreement, HEK293T cells expressing the mutated SAFA (Flag-Del-650-675) produced more type I-IFNs compared with wild-type SAFA upon VSV infection (Figure 6F and S6B-C). Besides, RGG domain deletion (Flag-Del-RGG) deprived SAFA of facilitating interferon production and neither the fragment 1-675 nor the fragment 675-825 could augment IFN-β activation (Figure 6F and S6C), indicating that the antiviral function of SAFA is depend on the integrity of the big N-terminal fragment and the C-terminal RGG domain. Further, we expressed these mutants into THP-1 cells (Figure S6D). Compared with that in the Flag-SAFA expressing cells, the synergistic effect of SAFA-induced interferon and ISGs production after VSV infection was increased in the Flag-Del-650-675 expressing cells and decreased in the Flag-Del-RGG expressing cells (Figure 6G and 6H). Consistently, the mutated SAFA (Flag-Del-650-675) suppressed the replication of VSV more effectively (Figure 6I). These data collectively demonstrated the importance of RNA binding ability of SAFA in antiviral immune response.

## Discussion

Chromatin accessibility plays a central role in regulation of gene expression. The innate immune system responses rapidly to invading pathogens, which immediately produces various cytokines to eliminate the infection. The presence of accurate and effective cytokines production that can resolve the infection without causing host pathology is pivotal for the host. We previously reported that the nuclear matrix protein SAFA surveils viral RNA and regulates antiviral gene expression (Cao et al., 2019a). However, how SAFA activates and regulates the expression and what determines the specificity of antiviral immune genes remains unknown. In the present study, we identified that SAFA regulated chromatin accessibility of antiviral gene and the SAFA-interacting RNA mapped the specific genomic sites. First, accumulating evidence has shown that SAFA is involved in regulation of chromatin structure from a compacted state to an active open state, indicating that SAFA showed important potential in chromatin remodeling (Fan et al., 2018; Nozawa et al., 2017). Second, our genome wide sequencing results by ATAC-seq, RNA-seq and ChIP-seq in wild-type and SAFA deficient cells after VSV infection showed that SAFA is essential for chromatin accessibility, gene expression and enhancers/super-enhancers activation of antiviral genes (Figure 1 and 2). Third, in the interphase, SAFA remodels chromatin structure through oligomerization with chromatin-associated RNAs (Nozawa et al., 2017). Our results suggest that the RNA binding ability of SAFA is also indispensable for its function in that the RGG domain depletion deprived its role in regulating chromatin accessibility after viral infection (Figure 3). Intriguingly, VSV infection induced cleavage of SAFA that removed the RGG domain (Figure 6), indicating the importance of RNA-binding ability of SAFA in antiviral response.

Modulation of chromatin accessibility determines which gene is to be transcribed and therefore, chromatin modulation determination during viral infection is critical (Klemm et al., 2019; Tsompana and Buck, 2014). Our results suggested that SAFA mediates the modulation of anti-viral chromatin accessibility and this process is dependent on its RNA-binding ability (Figure 3). Further RIP-seq results showed that the RNAs interacting with SAFA after viral infection are mainly antiviral related (Figure 4), and knockdown of these RNAs impaired the accessibility of specific genomic sites (Figure 5). These results indicate that the RNA product during viral infection mediated the accessibility of related genes. Rigorous regulation of cytokines production is a crucial cellular process, in which different kinds and levels of cytokines are produced at different stages of infection. RNA products are diverse and short-lived and reflect the transcriptional program directly, showing great potential in regulation of biological processes (Morris and Mattick, 2014; Roden and Gladfelter, 2021). There are growing evidence suggesting that RNAs play important roles in regulation of chromatin accessibility at defined genomic loci (Caudron-Herger and Rippe, 2012; Dong et al., 2020; Huo et al., 2020; Mousavi et al., 2013). Moreover, it has been reported that RNA product provides feedback on transcription via regulation of electrostatic interactions in transcriptional condensates (Henninger et al., 2021). SAFA protein showed multivalent interactions potential that it undergoes oligomerization after binding to RNAs during viral infection, indicating that condensates formation may occur during the activation of SAFA (Lin and Cao, 2020).

To escape the inhibitory effects of host immune system, viruses have evolved various mechanisms to dampen the immune response. During DNA virus infection, inflammatory caspases cleave cGAS at the N-terminal that renders its activity in facilitating type I interferons production (Wang et al., 2017). Cleavage of hnRNP-M is a general strategy utilized by picornaviruses to facilitate viral replication (Jagdeo et al., 2015). Here we report that VSV cleaved both human and mice SAFA protein, resulting in RGG domain depletion from SAFA (Figure 6).

Extranuclear signaling pathways were generally critical for anti-viral signal transduction. These signals eventually converge in the nucleus. SAFA, which predominantly localized in the nucleus, oligomerized with the generated RNA product and initiated and maintained the openness of corresponding genes. ATAC-qPCR results showed that the accessibility of antiviral immune genes was determined by the RNA product interacted with SAFA in an extranuclear signaling pathway dependent way (Figure S5B). Thus, intranuclear and extranuclear signaling pathways cooperate and form a transcriptionally responsive mesh that remodels chromosome structures and facilitates anti-viral innate immunity.

Taken together, our results provide insights into how SAFA and RNAs collaborate to reprogram chromatin modulation specificity and accessibility to regulate antiviral gene expression.

## Acknowledgments

We thank Xiong Ji (Peking University, Beijing) for help with Chromatin Immunoprecipitation Sequencing (ChIP-seq) and Transposase-Accessible Chromatin with high throughput sequencing (ATAC-seq). This work was supported by the National Key Research and Development Program of China (2016YFA0500300; 2020YFA0707800), the National Natural Science Foundation of China (31570891; 31872736; 32022028; 81991505; 32000113), Peking University Clinical + X (PKU2020LCXQ009) and the Peking University Medicine Fund (PKU2020LCXQ009).

## Author Contributions

F.Y. designed the study and revised the paper. L.C., YF.L., YJ.L., XF.G., S.L., M.Z., Y.Z., wrote the paper, performed experiments and analyzed the data. X.J. provided technical support and contributed to imaging. X.G., F.G., Q.Z. provided expertise.

## Declaration of Interests

The authors declare no competing interests.

## STAR★Methods

### Key Resources Table

| REAGENT or RESOURCE | SOURCE | IDENTIFIER |
| --- | --- | --- |
| **Antibodies** |  |  |
| Mouse Anti-hnRNP U (clone 3G6) antibody | Santa cruz | Cat# sc-32315, RRID:AB_627741 |
| Rabbit Anti-HA antibody | Sigma-Aldrich | Cat# H3663, RRID: AB_262051 |
| Rabbit Anti-Histone H3 antibody - ChIP Grade (acetyl K27) | Abcam | Cat# Ab4729, RRID:AB_2118291 |
| Rabbit Anti-Tubulin antibody | Affinity | Cat# AF7011 |
| Mouse anti-Histone H3 antibody | MBL International | Cat# MABI0301, RRID: AB_11142498 |
| Rabbit Anti-DDDDK-tag antibody (anti-Flag) | MBL | Cat# PM020, RRID:AB_591224 |
| Mouse Anti-Lamin B1 (clone 3C10G12) antibody | Proteintech | Cat# 66095-1-Ig, RRID:AB_11232208 |
| **Chemicals, Peptides, and Recombinant Proteins** |  |  |
| Protease inhibitor cocktail | TargetMol | Cat# B14001 |
| Pierce Protein A/G Agarose | Thermo | Cat#20422 |
| Lipofectamine 3000 Reagent | ThermoFisher | Cat#L3000015 |
| **Virus** |  |  |
| VSV (Vesicular Stomatitis Virus, Indiana strain) | J. Rose (Yale Univ) | N/A |
| **Experimental Models: Cell lines** |  |  |
| Human: 293T | ATCC | Cat#ATCC-CRL-3216 |
| Human: 2fTGH | Cao et al., 2019 | N/A |
| Human:THP1 | Zhengfan J. (Peking) | N/A |
| **Recombinant DNA** |  |  |
| Plasmid: Pcmv7.1 FLAG-SAFA | Cao et al., 2019 | N/A |
| Plasmid: Pcmv7.1 3xHA-SAFA | Cao et al., 2019 | N/A |
| Plasmid: Pcmv7.1 3xHA-Del650-675 | This paper | N/A |
| Plasmid: Pcmv7.1 3xHA- Del675-700 | This paper | N/A |
| Plasmid: Pcmv7.1 FLAG-Del650-675 | This paper | N/A |
| Plasmid: Pcmv7.1 FLAG-Del RGG | Cao et al., 2019 | N/A |
| Plasmid: Pcmv7.1 FLAG-1-675 | This paper | N/A |
| Plasmid: Pcmv7.1 FLAG-675-825 | This paper | N/A |
| Plasmid: Plvx FLAG-SAFA | This paper | N/A |
| Plasmid: Plvx FLAG-Del 650-675 | This paper | N/A |
| Plasmid: Plvx FLAG-Del RGG | This paper | N/A |
| pCMV-VSV-G | AddGene | Cat#8454 |
| psPAX2 | AddGene | Cat#12260 |
| pXR003: CasRx gRNA cloning backbone | AddGene | Cat#109053 |
| pXR001: EF1a-CasRx-2A-EGFP | AddGene | #109049 |
| Human IFNB1-Promoter-Luci: | Cao et al., 2019 | N/A |

### Lead Contact and Materials Availability

*Further information and requests for reagents may be directed to, and will be fulfilled by the corresponding author Fuping You (*.).

### Experimental Model and Subject Details

#### Cells

THP-1 cell was a gift from Zhengfan Jiang (Peking University). 2fTGH-ISRE cell (human fibrosarcoma cell expressing an ISRE driven luciferase reporter) was generated by stabilizing ISRE-luciferase plasmid in 2fTGH cell. Isolation of BMDM (bone-marrow derived macrophages) was performed as described (Cao et al., 2019a). *SAFA*^-/-^, *MAVS*^-/-^, *IRF3*^-/-^ THP-1 cells were constructed by CRISPR-Cas9 system as previously reported (Cao et al., 2019a). Cells were cultured in Dulbecco’s Modified Eagle’s Medium (DMEM) supplemented with 10% FBS, 100 U/mL Penicillin-Streptomycin. Cells were negative for mycoplasma.

#### Constructs

Expression constructs generated for this study were prepared by standard molecular biology techniques and coding sequences were entirely verified. All the deletions and mutants were constructed by standard molecular biology technique. Each construct was confirmed by sequencing.

### Method Details

#### Type I IFN Bioassay

Type I IFN bioassay was performed as previously reported (Cao et al., 2019a). Type I IFNs in human cell culture medium were quantified using a 2fTGH-ISRE cell line stably expressing an ISRE-Luci reporter. In brief, 200 mL of culture medium was incubated with confluent 2fGTH-ISRE-Luci cells (24-well plate) for 6 hours. Cells were lysed in passive lysis buffer and subjected to luciferase quantification (Promega). A serial dilution of human IFNb was included as standards.

#### Luciferase Reporter Assay

Luciferase reporter assay was performed as previously reported (Cao et al., 2019b). HEK293T cells seeded on 24-well plates were transiently transfected with 50 ng of the luciferase reporter plasmid together with a total of equal amount of indicated expression plasmids or empty control (EV) plasmid. As an internal control, 10 ng pRL-TK was transfected simultaneously. Reporter gene activity was analyzed using the Dual-Luciferase Reporter 1000 Assay System (Promega) and measured with a TD-20/20 Luminometer (Turner Designs) according to the manufacturers’ instructions.

#### Plaque Assays

Viral titers from the cell culture medium were determined by plaque-forming assays as previously described (35). Briefly, virus-containing medium was serially diluted and then added to confluent Vero cells. After incubation for 1 hour, supernatants were removed, cells were washed with PBS, and culture medium containing 2% (wt/vol) methylcellulose was overlaid for 24 hours. Then cells were fixed for 30 minutes with 0.5% (vol/vol) glutaraldehyde and then stained with 1% (wt/vol) crystal violet dissolved in 70% ethanol for 30 minutes. After washing twice with ddH_2_O, plaques were counted, and average counts were multiplied by the dilution factor to determine the viral titer as plaque-forming units per milliliter.

#### Western blotting

Cells were harvested and lysed with Pierce lysis buffer (25 mM Tris·HCl, pH 7.4, 150 mM NaCl, 1 mM EDTA, 1% NP-40, 5% β-Mercaptoethanol) with the protease inhibitor cocktail (Roche) on ice for 30 minutes. Supernatants were collected by centrifugation at 12,000 rpm for 10 minutes at 4°C. Cell lysates were boiled with loading buffer. Each protein sample was loaded onto 8 % SDS-PAGE. After electrophoresis, proteins were transferred to the nitrocellulose membrane (Millipore). The membrane was blocked with 5% milk (in PBST) for 1 hour, and incubated sequentially with primary and HRP-coupled secondary antibodies. After being washed with PBST for 3 times, the membranes were visualized by enhanced chemiluminescence (Millipore).

#### Coomassie brilliant blue staining

Samples pulled down with SAFA antibody were anaylzed with SDS-PAGE. After staining with Coomassie brilliant blue R-250, the target bands on the PAGE gel were visualized and excised for mass spectrometry.

#### RNA knockdown

RNA knockdown was performed by CRISPR-Cas13d system. Specific CRISPR RNAs (crRNAs) were annealed and ligated into CasRx gRNA cloning backnone (addgene, #109053). CrRNA plasmids (2 ug) and plasmids coding CasRx (addgene, #109049) were transfected into HEK293T cells together (6-well plate). The medium was changed to fresh DMEM containing 10% FBS at 6 hours post transfection. After transfection for 48 hours, GFP-highly positive cells were sorted by using fluorescence-activated cell sorting (FACS) and used for further experiment. The crRNAs used were listed in the table S1.

#### Quantitative real-time PCR (qRT-PCR)

Total RNA was isolated using the RNA simple Total RNA kit (TIANGEN). 1 ug RNA was reverse transcribed using a FastKing RT Kit (TIANGEN). Levels of the indicated genes were analyzed by qRT-PCR amplified using SYBR Green (Transgene). Data shown are the relative abundance of the indicated mRNA normalized to *Actin*. The primers used were listed in the table S2.

#### ATAC-seq

Pellet 50,000 viable sample cells at 500 RCF at 4°C for 5 min. Aspirate all supernatant. Add 50 μL cold ATAC-Resuspension Buffer (RSB) containing 0.1% NP40, 0.1% Tween-20, and 0.01% Digitonin into the cell pellet and pipette up and down 3 times. Incubate on ice for 3 minutes. Wash out lysis with 1 mL cold ATAC-RSB containing 0.1% Tween-20 but no NP40 or digitonin and invert tube 3 times to mix. Pellet nuclei at 500 RCF for 10 min at 4°C. Aspirate all supernatant. Resuspend cell pellet in 50 μL of transposition mixture (25 μL 2x TD buffer, 2.5 μL transposase (100 nM final), 16.5 μL PBS, 0.5 μL 1% digitonin, 0.5 μL 10% Tween-20, 5 μL H_2_O) by pipetting up and down 6 times. Incubate reaction at 37°C for 30 minutes. Afterward, the DNA was purified with Magen DNA purify kit and amplified with primers containing barcodes by using the TruePrep DNA Library Prep Kit (TD501-01). All libraries were adapted for sequencing.

#### ATAC-qPCR

The ATAC libraries were adapted for qRT-PCR with specific primers. The primers used were listed in the table S3.

#### RNA-seq

Bulk RNA-sequencing (RNA-seq) was conducted as previously described (Cao et al., 2019a). The original data of the RNA-seq was uploaded to the GEO DataSets.

#### RIP-seq

RNA immunoprecipitation sequencing (RIP-seq) was conducted as previously described (Cao et al., 2019a). The original data of the RIP-seq was uploaded to the GEO DataSets.

#### ChIP-seq

Chromatin immunoprecipitation followed by sequencing (ChIP-seq) and data analysis were conducted as previously described (Cao et al., 2019a). The original data of the ChIP-seq was uploaded to the GEO DataSets.

#### Statistical Analysis

For all the bar graphs, data were expressed as means ± SEM. Prism 8 software (graphic software) was used for charts, and statistical analyses. Differences in means were considered statistically significant at *p* < 0.05. Significance levels are: * *p* < 0.05; ** *p* < 0.01; *** *p* < 0.001; **** *p* < 0.0001; NS., non-significant.

#### Data availability

ATAC-seq, ChIP-seq, RNA-seq and Rip-seq relevant data are available at the DRYAD database: https://doi.org/10.5061/dryad.0rxwdbs0w.

#### Tables

Table S1: CrRNA sequence

Table S2. Primers for qRT-PCR

Table S3. Primers for RT-PCR and ATAC-qRT-PCR

---

**Figure S1.**
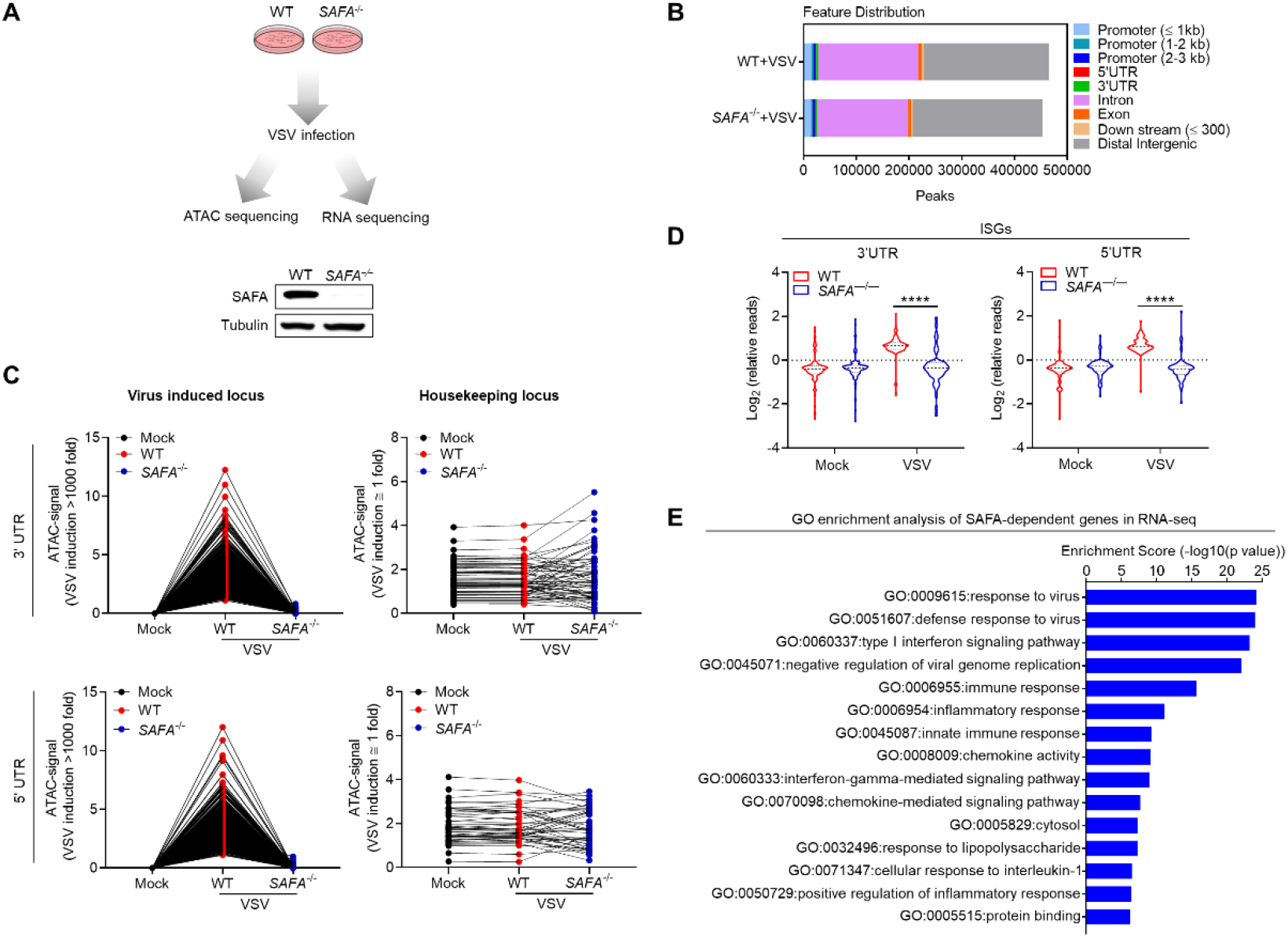
SAFA deficiency decreased the chromatin accessibility of antiviral immune genes. (A) Models depicting the ATAC-seq and RNA-seq in Wild-type (WT) and *SAFA*^-/-^ THP-1 cells with VSV infection for 6 hours (upper), and immunoblotting results showing the knockout of SAFA in THP-1 cells (lower). (B) Feature distribution of ATAC-seq profile after VSV infection in WT and *SAFA*^-/-^ THP-1 cells. (C) Line graph showing SAFA in regulation of VSV induced accessible locus and insensitive locus. (D) Violin graph showing ISGs affected by SAFA depletion in ATAC-seq. (E) GO term enrichment analysis of genes significantly affected by SAFA depletion in RNA-seq. \*\*\*\**p* < 0.0001 (Student’s t test; D). Data were pooled from two independent experiments (B, C and E).

**Figure S2.**
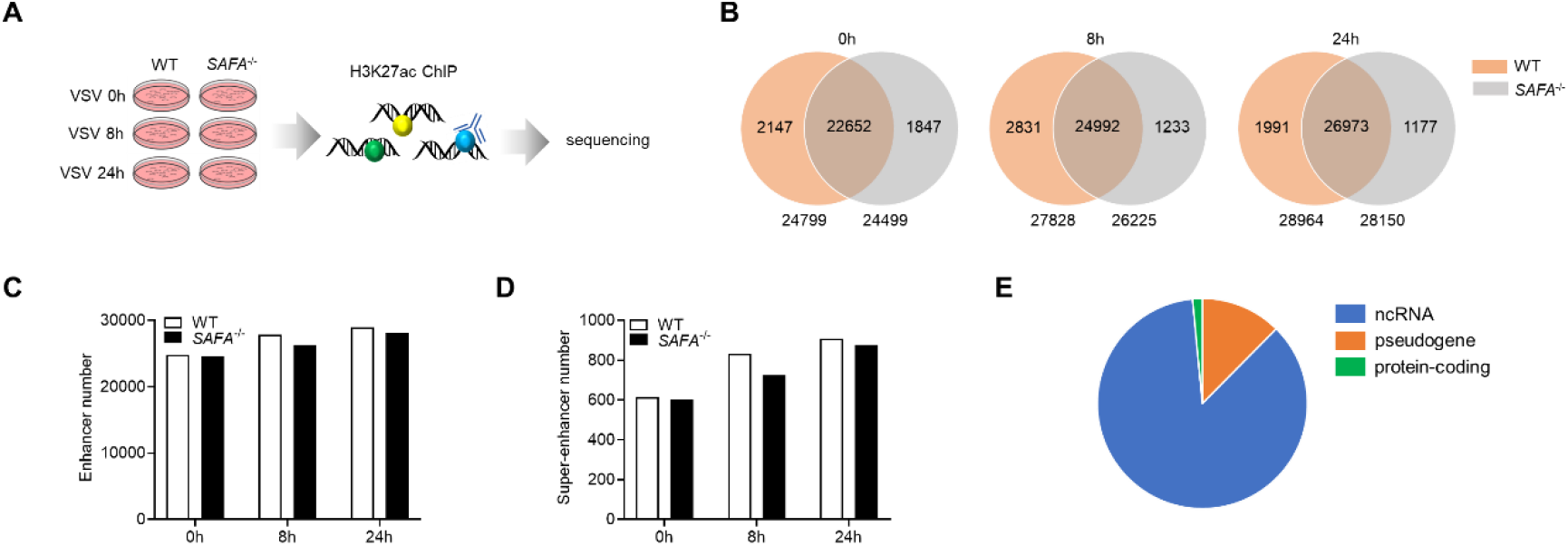
SAFA deficiency decreased the activation of antiviral immune genes. (A) Models depicting the ChIP-seq assay of H3K27ac in Wild-type (WT) and *SAFA*^-/-^ THP-1 cells with VSV infection for 6 hours. (B) Venn diagram showing amounts of enhancers in WT and SAFA^-/-^ THP-1 cells with VSV infection. (C) Histogram diagram showing amounts of enhancers in WT and SAFA^-/-^ THP-1 cells with VSV infection. (D) Histogram diagram showing amounts of super-enhancers in WT and SAFA^-/-^ THP-1 cells with VSV infection. (E) Pie graph showing distribution of super-enhancer-driven genes. Data were pooled from two independent experiments (B-E).

**Figure S3.**
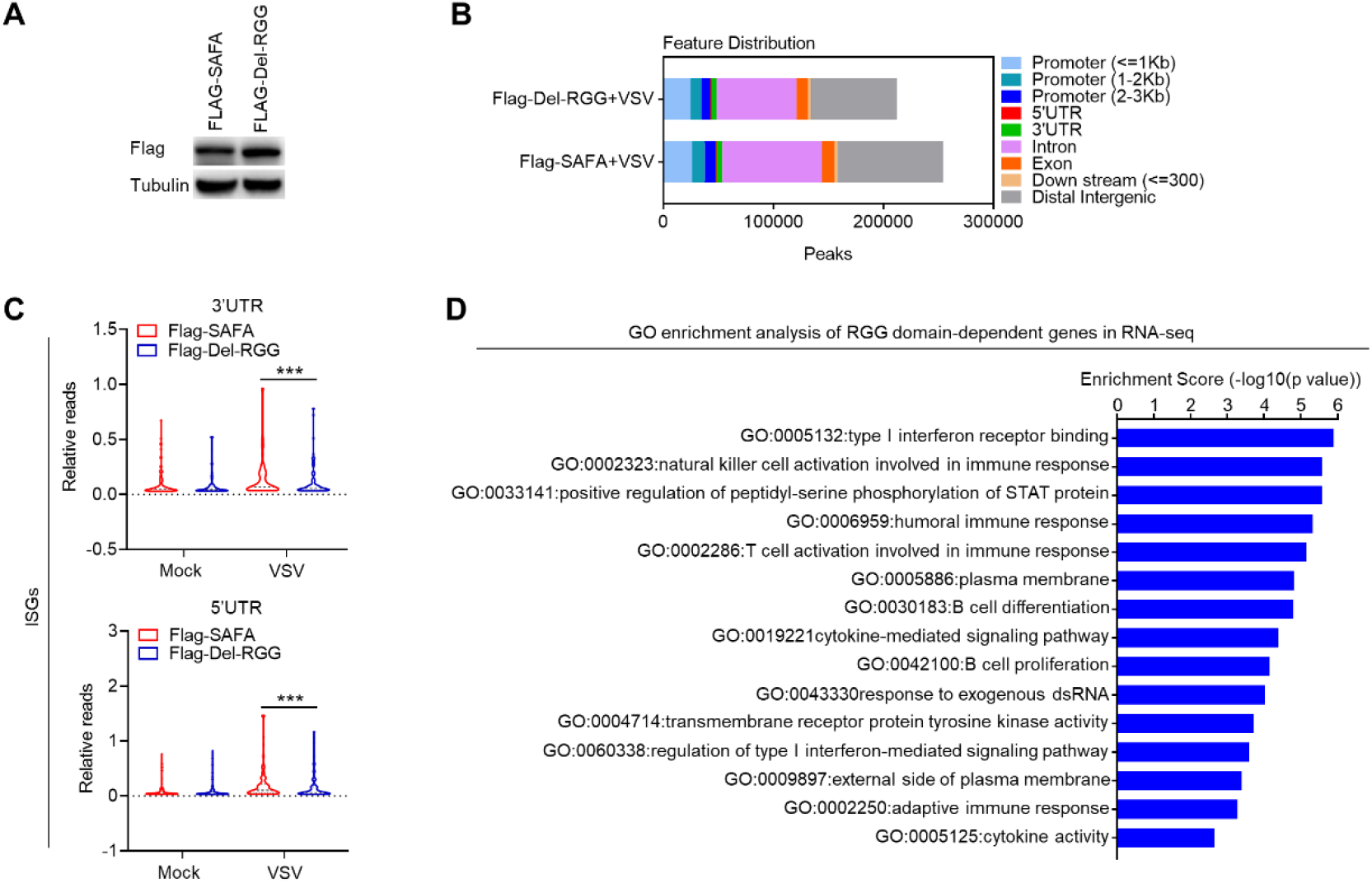
RNA binding activity of SAFA is critical for increasing the accessibility of anti-viral chromatin. (A) Immunoblotting results showing the expression of Flag-SAFA and Flag-Del-RGG in *SAFA*^-/-^ THP-1 cells (B) Feature distribution of ATAC-seq profile after VSV infection (C) Violin graph showing ISGs affected by RGG domain depletion in ATAC-seq (D) GO term enrichment analysis of genes significantly affected by RGG domain depletion in RNA-seq \*\*\**p* < 0.001 (Student’s t test; C). Data were pooled from two independent experiments (B and D). Data were representative of two independent experiments (A).

**Figure S4.**
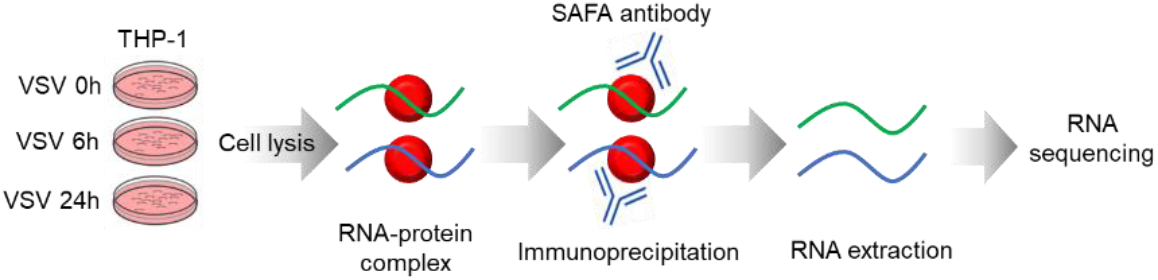
SAFA interacted with antiviral related RNAs in a time-dependent manner during viral infection. Models depicting the RIP-seq assay of SAFA in THP-1 cells with VSV infection for 6 or 24 hours.

**Figure S5.**
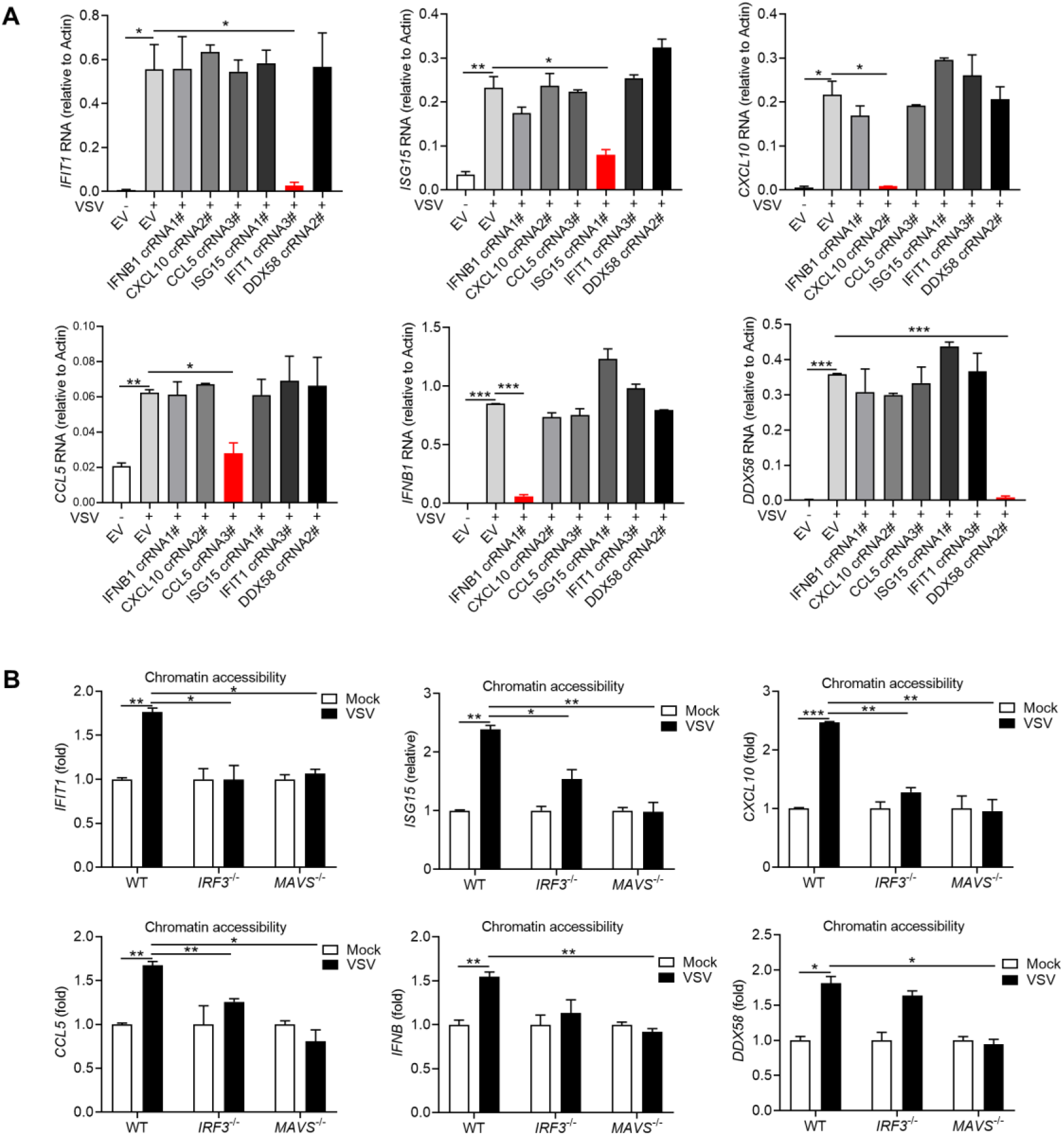
SAFA-interacting RNA mediated specific chromatin remodeling in an extranuclear pathway dependent manner. (A) Histogram showing the RNA expression with indicated crRNA transfection for 48 hours and with or without VSV infection for 18 hours (B) ATAC-qPCR results showing the chromatin accessibility of indicated genes after VSV infection for 18 hours in WT, *IRF3*^-/-^ and *MAVS*^-/-^ THP-1 cells. \**p* < 0.05, \*\**p* < 0.01, \*\*\**p* < 0.001 (Student’s t test). Data were pooled from three independent experiments. Error bars, SEM. n = 3 cultures.

**Figure S6.**
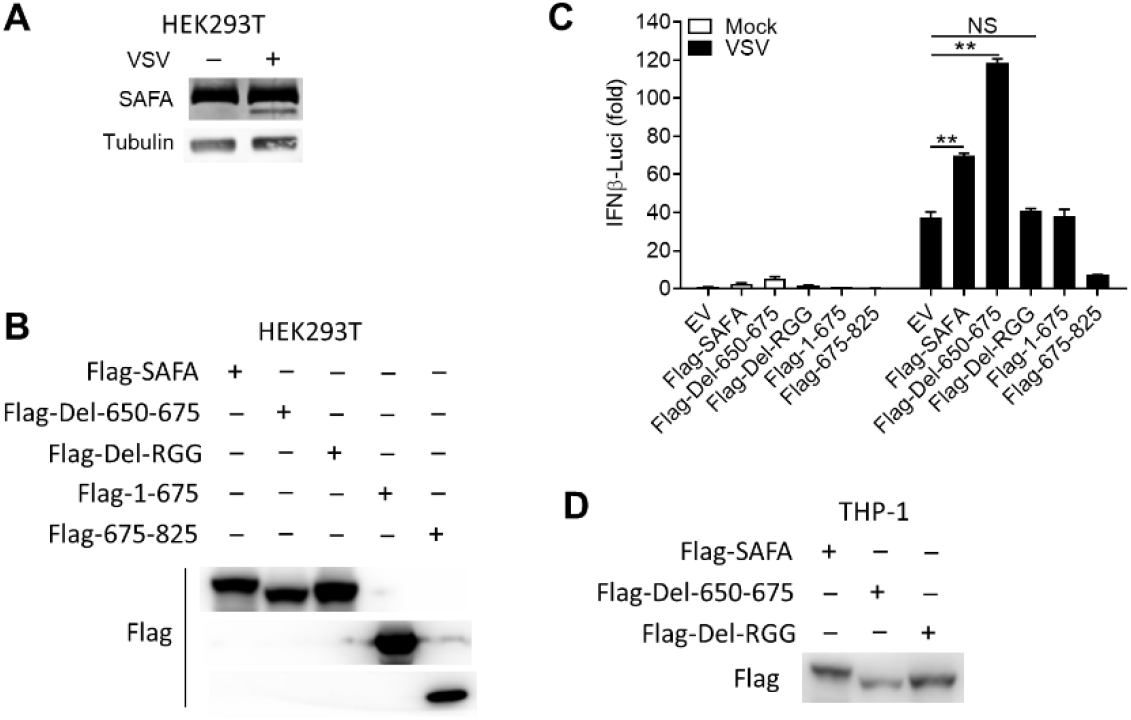
Virus-mediated cleavage separates the RNA-binding domain from SAFA. (A) HEK293T cells were infected with VSV for 4 hours, and the indicated protein were detected by immunoblotting. (B) HEK293T cells were transfected with indicated plasmids, and the expression level of these plasmids were detected by immunoblotting (C) Luciferase activity of IFNβ in HEK293T cells expressing IFNβ–Luc plasmid together with either an empty vector or indicated plasmids, after 24 hours infected with VSV for 24 hours. (D) THP-1 mutants were generated by overexpressing indicated lentivirus plasmids, and the expression level of these plasmids were detected by immunoblotting. \*\**P* < 0.01 (Student’s t-test). Data were representative of three independent experiments (A, B and D). Data were pooled from 3 independent experiments (C). Error bars, SEM. n = 3 cultures.

**Table S1:**
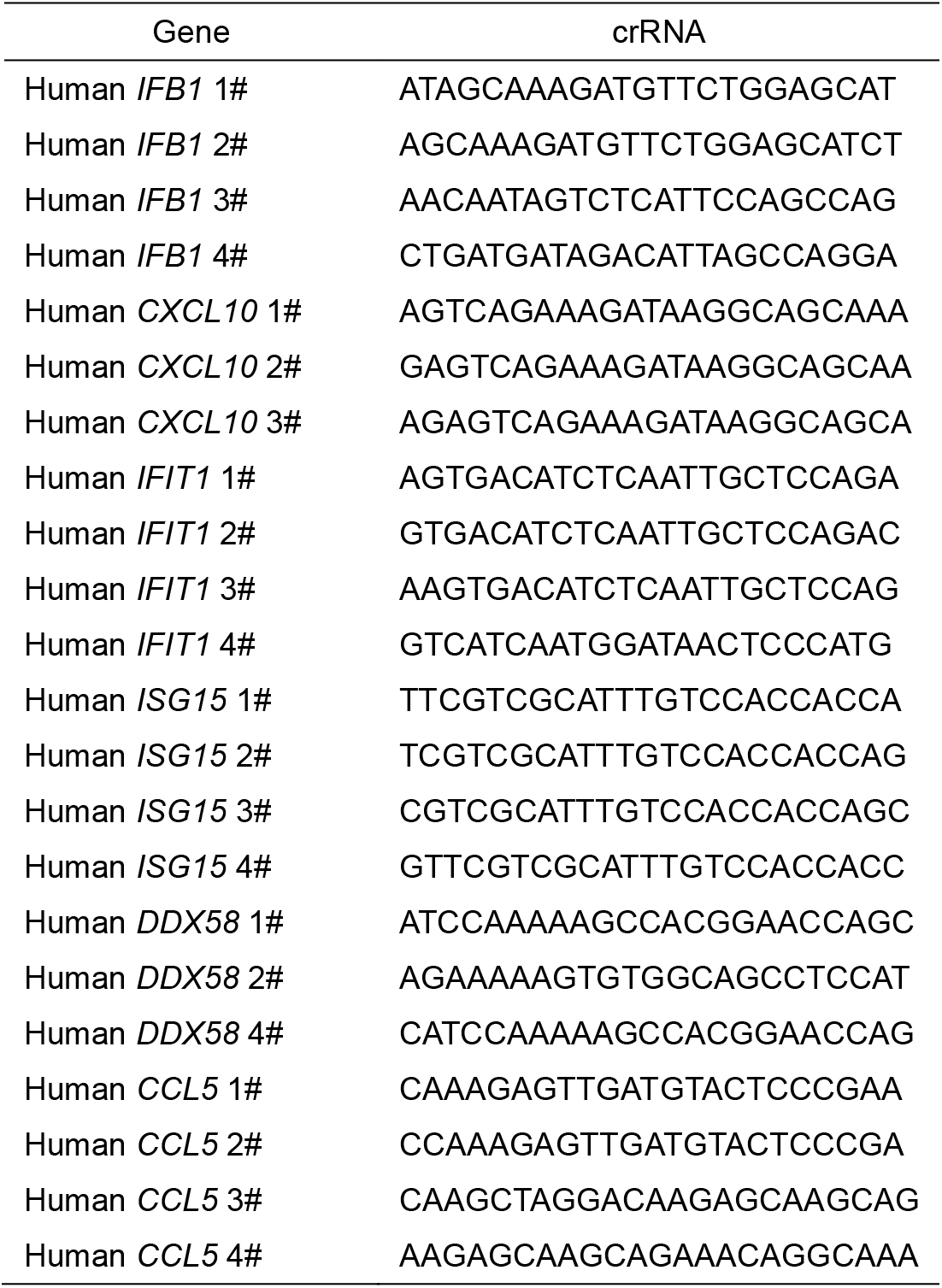
CrRNA sequence

**Table S2.**
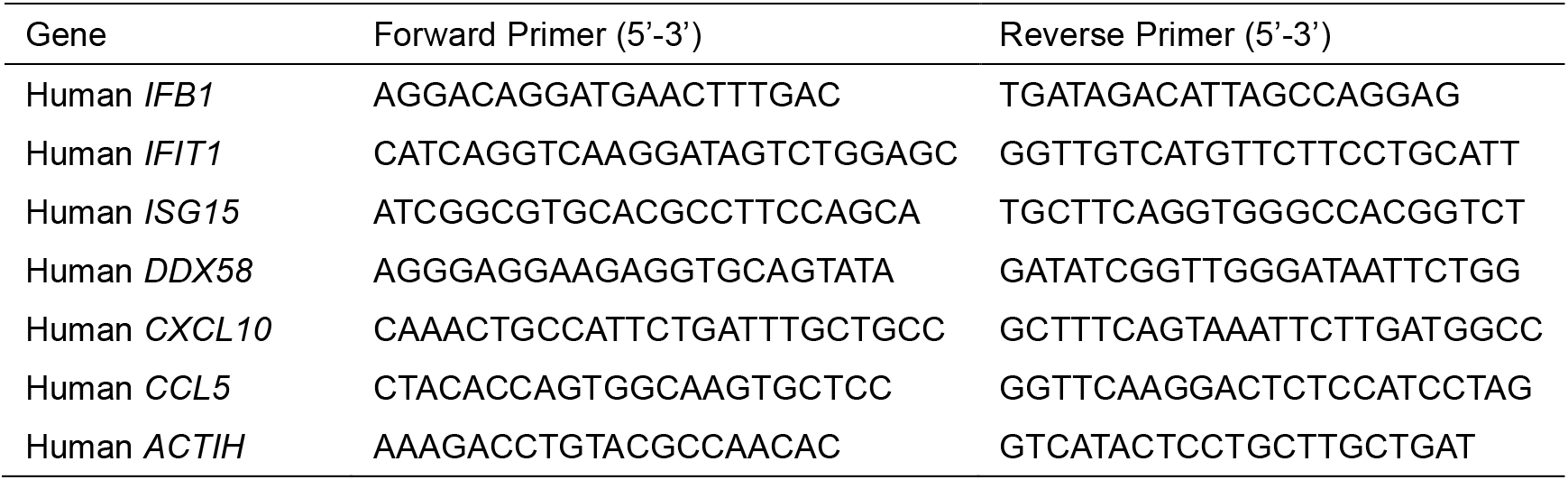
Primers for qRT-PCR

**Table S3:**
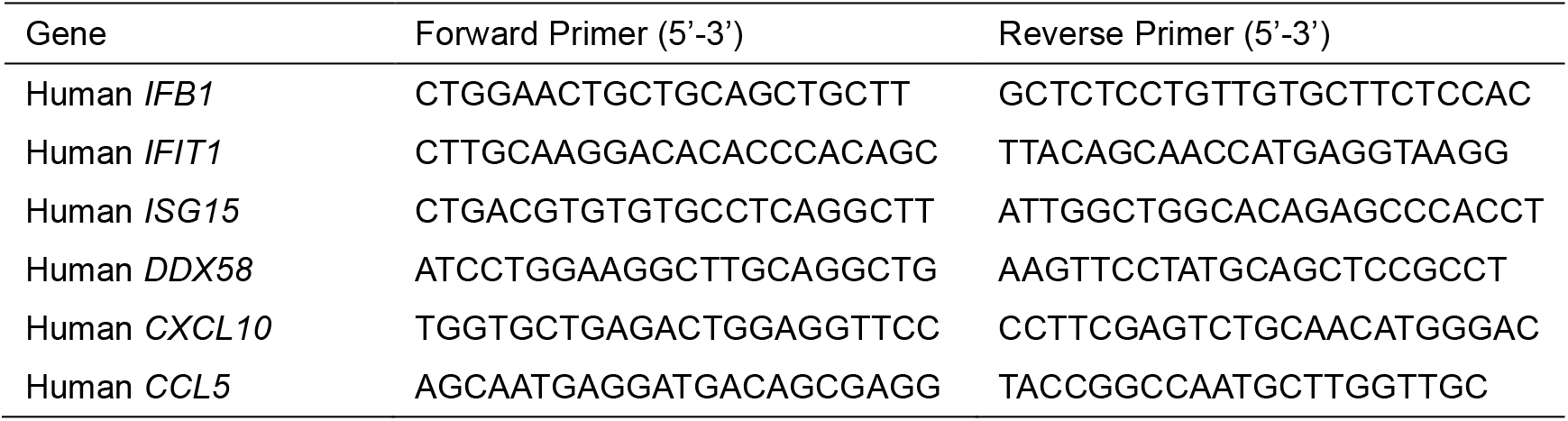
Primers for ATAC-qRT-PCR

## Notes

### Competing Interest Statement

The authors have declared no competing interest.

